# A Hallmark-Integrated, Agent-Based Framework for Intratumor Heterogeneity in Melanoma Evolution

**DOI:** 10.64898/2026.01.16.699757

**Authors:** Khola Jamshad, Trachette L. Jackson

## Abstract

Advances in multiregion sequencing have revealed extensive intratumor heterogeneity (ITH) - the presence of genetically distinct subclones within a single tumor. ITH profoundly influences tumor behavior, including well-accepted hallmarks of cancer such as sustained proliferation, resistance to apoptosis, and immune evasion, and the emergence of therapeutic resistance. In this work, we introduce a hallmark-integrated branching evolution process agent-based model (BEP-HI) to study the emergence of intratumor heterogeneity (ITH) in melanoma under coupled genetic, immune, and spatial selection pressures. Using this model, we identify three distinct evolutionary ITH modes and demonstrate a mechanistic decoupling between tumor growth kinetics and heterogeneity. We further show that immune recruitment exerts the strongest influence on ITH through nonlinear immune-editing feedback, while motility of melanoma cells shapes complex morphologies in different ITH modes as observed in clinical SSM tumors. This work provides a biologically grounded and adaptable computational framework for exploring how hallmark interactions shape ITH evolution and for generating virtual tumor cohorts that link genetic diversity with tumor behavior, morphology, and treatment resistance.

## 1 Introduction

Understanding how tumors evolve is one of the most urgent and complex challenges in modern oncology. This complexity arises because of a process of evolution playing out at the cellular level, driven by the accumulation of genetic mutations and shaped by interactions with the tumor microenvironment (TME). Advances in multiregion sequencing have revealed extensive intratumor heterogeneity (ITH) - the presence of genetically distinct subclones within a single tumor. These findings challenge earlier models that assumed tumor progression followed a linear, stepwise accumulation of mutations. Instead, a branching evolution process (BEP) model has gained prominence, in which multiple subclonal lineages evolve in parallel from a common ancestor, resulting in spatial and temporal heterogeneity. Mutations that occur early in this process and are therefore common in nearly all of the clones are known as founder or trunk mutations and those that occur later on are known as progressor or branch mutations (Davis et al. 2017). Mutations in driver genes confer fitness advantages to the cell and drive tumor evolution whereas passenger mutations do not confer fitness advantages but occur in a cell that coincidentally or subsequently acquires a driver mutation, and takes these passenger mutations along with it through the tumor evolution process (Bozic et al. 2010). ITH profoundly influences tumor behavior, including well-accepted hallmarks of cancer such as sustained proliferation, resistance to apoptosis, and immune evasion. It also underlies the emergence of therapeutic resistance, as distinct subpopulations may harbor or acquire mutations that confer survival advantages under treatment pressure. Therefore, ITH plays a significant role in shaping the tumor landscape.

Melanoma, a skin cancer with the highest somatic mutation burden among solid tumors, provides a powerful system for examining how ITH shapes tumor evolution. Targeted sequencing and gene-expression studies have revealed extensive heterogeneity within individual melanomas (Harbst et al. 2014), and multiple analyses show that even a single metastatic lesion may arise from several distinct subpopulations within the primary tumor (Sanborn et al. 2015). Such diversity has profound clinical implications. For example, resistance to targeted therapy or immunotherapy often reflects multiple mechanisms operating simultaneously within a single tumor, including both pre-existing resistant clones and new mutations acquired during treatment (Kemper et al. 2015). Understanding how these processes interact is essential for predicting tumor trajectories and designing strategies to prevent or overcome resistance.

Although melanoma is highly immunogenic, it has a remarkable capacity to evade immune surveillance, further amplifying ITH. Melanoma cells can reduce antigenicity or immunogenicity by downregulating tumor-associated antigens, losing MHC class I expression, or disrupting antigen-presentation pathways, thereby limiting recognition by cytotoxic T cells (CTLs) (Eddy et al. 2021). Under sustained immune pressure, distinct immune-resistant subclones may emerge and remodel the tumor microenvironment, limiting immune infiltration and function. This interplay between genetic diversification and immune editing contributes to immunological ITH, spatially compartmentalized clonal evolution, and variable therapeutic responses across—and even within—patients. These intricate dynamics motivate the need for computational approaches that can capture spatial and genetic complexity in tumor dynamics.

Agent-based models (ABMs) are especially well-suited for capturing these emergent dynamics by explicitly modeling individual cell behavior, local interactions, and environmental context. Waclaw et al. used a 3D lattice-based model to simulate the spatial spread of mutations during tumor growth (Waclaw et al. 2015) and found that spatial constraints can suppress clonal mixing, resulting in low effective heterogeneity despite high mutation rates. Similarly, Gerlee and Anderson focused on trait evolution, simulating how proliferation and motility evolve in spatially structured tumors to show that restrictions in spatial arrangement of cells slow selection and preserve diversity (Gerlee and Anderson 2015). While these models establish foundational insights into spatial evolution, they do not account for immune interactions or therapeutic selection pressures.

To understand how melanoma tumors evolve, it is critical to develop models that accurately reflect the underlying biological and genetic complexity of the disease. Melanoma is driven by a well-characterized set of mutations in genes such as BRAF, NRAS, and KIT etc., each of which contributes differently to cell proliferation, survival, and immune evasion. These drivers not only define molecular subtypes but also influence spatial growth patterns and treatment response. However, existing models often oversimplify these dynamics, failing to capture how genetically distinct subclones compete, cooperate, or adapt within the tumor microenvironment. A data-driven ABM provides a framework to directly incorporate driver gene related data and eventually patient-derived data, such as mutation frequencies, spatial gene expression, and cell phenotyping, into simulations that reflect biologically plausible tumor behavior. By grounding model parameters and cell decision rules in empirical data, such ABMs can better reproduce observed patterns of ITH and therapy resistance. Developing a comprehensive understanding of how genetic heterogeneity emerges and evolves, particularly in relation to spatial organization within the tumor and its interactions with the immune system, is critical for developing more effective, personalized therapies.

In this work, we introduce a hallmark-integrated branching evolution process agent-based model (BEP-HI) to study the emergence of intratumor heterogeneity (ITH) in melanoma under coupled genetic, immune, and spatial selection pressures. Unlike prior BEP and spatial tumor models that focus primarily on proliferative fitness, our framework explicitly integrates multiple cancer hallmarks — genetic instability, survival, immune evasion, and proliferation — mapped onto melanoma-relevant driver mutations within a spatially resolved tumor–immune microenvironment. Using this model, we identify three distinct evolutionary ITH regimes and demonstrate a mechanistic decoupling between tumor growth kinetics and heterogeneity: driver strength governs time to detectable tumor size, whereas mutation rate and immune recruitment control transitions between clonal, subclonal, and fractal heterogeneity. We further show that immune recruitment exerts the strongest influence on ITH through nonlinear immune-editing feedback, highlighting how immune pressure can reshape both genetic diversity and spatial organization. Together, this work provides a biologically grounded computational framework for exploring how hallmark interactions shape melanoma evolution and for generating virtual tumor cohorts that link genetic diversity, immune context, and tumor morphology.

## 2 Methods

### 2.1 Model development

Our work builds upon and significantly extends an existing agent-based branching evolution process (BEP) model (Niida et al. 2015). Their BEP model provided valuable insights into clonal dynamics by incorporating a single hallmark of cancer: increased proliferative capacity. While foundational, this focus limits the model’s ability to reflect real evolution which occurs through interaction across several hallmarks of cancer, often shaped by immune interactions and genomic instability. To address these limitations, we introduce a two-dimensional BEP-HI (Branching Evolutionary Process with Hallmark Integration) model, a novel framework that incorporates not only increased proliferation but also enhanced survival, immune evasion, and the enabling characteristic of genetic instability. Rather than a simple extension of the original model, BEP-HI represents a conceptual shift—moving from a one-dimensional fitness landscape to a multidimensional evolutionary space reflective of hallmark plasticity and selection pressures. We model BRAF-associated superficially spreading melanoma (SSM), the most common melanoma type, which starts as an irregularly-shaped flat lesion that grows radially (Druskovich et al. 2024). Later, we extend the model (BEP-HIM) by introducing cell motility to melanoma cells with the BRAF mutation.

In BEP-HI, there are two agents - melanocytes and immune cells. Melanocytes have a genome consisting of 50 genes of which seven are driver genes, and an anti-genicity associated with them. Each gene is represented as a binary value 0 (wild) or 1 (mutated). A melanocyte is considered to be a melanoma tumor cell (MTC) only when it acquires a driver mutation, it is otherwise a normal cell (NC). We initialize with 9 NCs (no mutations) and 1 MTC that has only one mutated founder driver gene (BRAF) for modeling SSM. The tumor grows in a two-dimensional lattice until it reaches either a maximum tumor population *c_max_* or maximum time steps *t_max_*. New cells have the same genetic profile as their parent cell and are placed in the Moore neighborhood of the parent. The stochastic actions of melanocytes and immune cells are summarized in Figure 1. A melanocyte might undergo apoptosis with a probability *q*, otherwise it can proliferate at probability *p*. If the cell undergoes proliferation, each gene in its genome has the opportunity to mutate with probability *r*. At any time, the number of driver genes *d_i_* associated with each hallmark *i* are tracked where *i* ∈ {*r*: mutation (genetic instability), *p*: increased proliferation, *q*: increased survival, *c*: decreased antigenicity, *c*°: increased antigenicity}. Driver gene strength *f* acts as a scaling factor for the degree of impact for each driver gene. Then action probabilities are: *p* = *p*_0_10*^fdp^*, *q* = *q*_0_10^−^*^fdq^*, and *r* = *r*_0_10*^fdr^* where *i*_0_ are the base probabilities associated with the hallmark, either from literature or tuned for the model. Melanoma cells have an antigenicity value *a* = *a*_0_10*^f^*^(^*^dc^*^°−^*^dc^*^)^, where base antigenicity *a*_0_ ∈ [1, 2] and the effect of driver genes increasing or decreasing antigenicity (aiding immune evasion) is accounted for. We introduce an immune stimulatory factor (ISF), *F* (**x**) which is determined at grid point **x** by antigenicity of cells within limited radius of **x**. Immune cells are recruited from the TME to the tumor periphery with probability *s* by a Poisson process where the rate *s_T_* is given by a Hill function

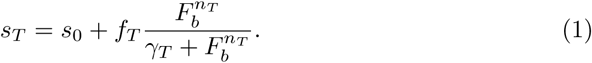

**Fig. 1:**
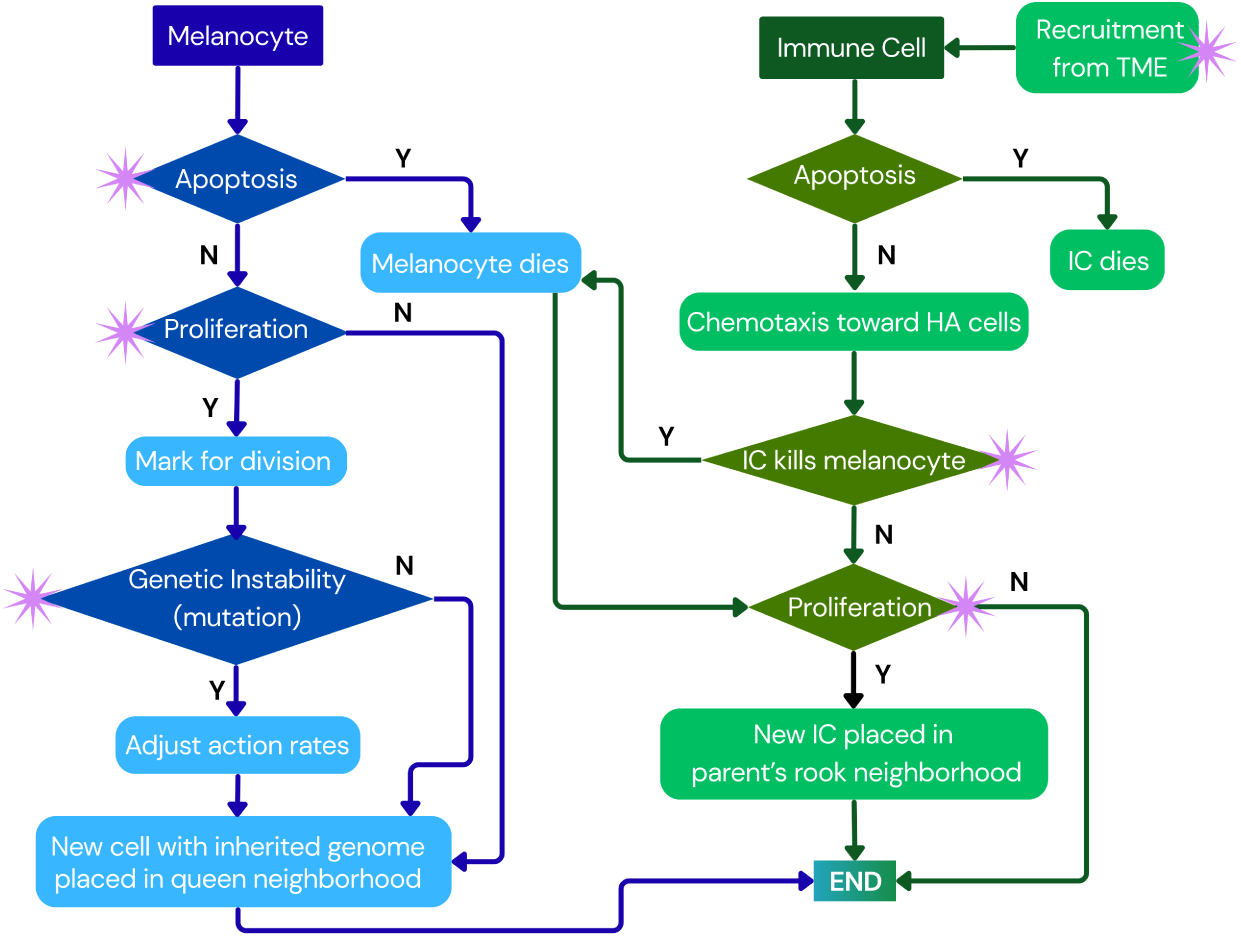
Schematic diagram of BEP-HI ABM for both melanocytes and immune cells, with asterisks indicating where action probabilities change depending on the driver mutations in the cell.

The base recruitment *s*_0_ *>* 0 allows initial CTL recruitment to the tumor site and we limit consideration to total ISF of border tumor cells (*F_b_*). Immune cells have a constant apoptosis probability. However, they proliferate (*p_i_*) and kill melanoma cells (*k*) in an ISF-dependent manner. Immune cells move by chemotaxis up the ISF gradient toward high antigenicity MTCs. Details on model implementation can be found in Appendix A with relevant parameters in Table 2.

### 2.2 Quantifying tumor heterogeneity and morphology

We visualize our tumor population such that variation in the color corresponds with genetic variation. To quantify the intratumor heterogeneity, we define the entropy of the population *ɛ*, which is a variance measure so that if we have a completely monoclonal population then *ɛ* = 0 and it increases with heterogeneity (Niida et al. 2015). We find the similarity matrix *S* from the *n* x *m* mutation profile matrix *M* of all melanocytes where *n* = 50 is genome size and there are *m* cells. Let *S_ij_* be the similarity between two cell genomes **g***^i^* and **g***^j^*, we then apply singular value decomposition to **S** to get a singular vector **s**.

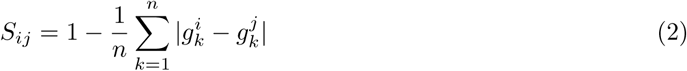

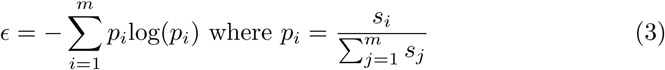

Additionally, we track the average mutation count per cell *β* to approximate the tumor mutational burden (TMB). We further look at surface fractal dimension (FD) analysis for simulated tumors. In our implementation, we cluster the tumor into subclones based on their driver gene mutational combination using k-means clustering, then apply the box counting method to find the *FD* for each cluster, and use their mean to represent the tumor ITH.

The morphology of a tumor plays a determining role in its invasive capability; irregularly shaped tumors are more likely to branch out and have greater opportunities for cells to detach and have access to the vascular system. In order to quantify the irregularity of our tumors, we look at their convexity measure 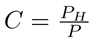 which compares the perimeter *P* of the tumor to the perimeter of the convex hull *P_H_* (smallest two dimensional polygon enclosing all MTCs). Lower values of *C* indicate irregular and highly infiltrative growth. We also consider the border fractal dimension (*FD_b_*) in which we generate a binary mask of the tumor and use that to determine its boundary, then use the box counting method. Higher *FD_b_* indicates more complex and irregular structures.

### 2.3 Sampling strategy

Given the importance of accurately capturing spatial heterogeneity in our tumor simulations, the ideal approach would involve analyzing the genetic data from the entire final cell population. However, because simulations typically generate populations exceeding 50,000 cells, this level of comprehensive sampling was computationally prohibitive. To balance computational feasibility with representational fidelity, we evaluate four sampling strategies, including simple random sampling (SRS), stratified sampling via CLARANS (CC), time systematic (TS), and space systematic (SS). Details on the implementation of each strategy can be found in Appendix B. The sample size was set at 10% of the population based on preliminary results indicating that simple random sampling (SRS) yields stable estimates of both *ɛ* (entropy) and *β* (mean mutation load) at this size. The mean error for each method is found with the true values of *ɛ* and *β* being calculated using the entire tumor cell population of 25,000 cells (3 runs per heterogeneity type).

### 2.4 Data integration of ITH in melanoma

Key oncogenes and tumor suppressor genes (TSGs) that undergo mutation in melanoma are listed in Table 1 in Appendix A along with the hallmarks they are associated with. These driver genes are selected based on the frequency of their occurence in melanoma and their contributions to heterogeneity. In melanomas, the somatic BRAF mutation rate is the highest among cancer types, occuring in about 50% of melanoma-affected patients. The constitutively activated BRAF affects the MAPK pathway thereby promoting tumour cell proliferation, survival, invasion, and evasion of the immune response. BRAF is most commonly associated with SSM tumors. BRAF, NRAS, and KIT are foundational melanoma mutations, and rarely overlap in a cell - this mutually exclusive behavior is built into the model.

**Table 1:** List of oncogenes and tumor suppressor genes that are mapped onto the driver genes in the ABM.

| Gene | Hallmarks |
| --- | --- |
| *BRAF*<sup>V600E</sup> | Growth, Survival, Immune Evasion, Genetic Instability |
| NRAS | Growth, Survival, Genetic Instability |
| KIT | Growth, Genetic Instability |
| MITF | Growth, Survival, Immune Evasion, Genetic Instability |
| CDKN2A | Growth |
| COX2 | Growth, Survival, Immune Evasion |
| PTEN | Growth, Survival, Immune Evasion |

### 2.5 Estimation of model parameters for SSM

We assume that a single melanoma cell is around 0.01 *mm* in diameter (Busam et al. 2001) and scale each ABM cell to 5 real cells. This results in our simulated tumors reaching a detectable size of about 6 *mm* in-vivo when the number of simulated cells reaches around *c_max_* = 50, 000 cells. The model is calibrated using driver gene strength *f* which impacts all hallmarks, and the enabling characteristic of genetic instability related base mutation rate *r*_0_. For calibration of *f*, we compared the time taken for SSM tumors to become detectable in-vivo with our BEP-HI simulation times to reach *c_max_* cells as *f* is varied. We further examine the growth rates of SSM tumors around time of detection and used it to find *r*_0_ values that work for our selected *f* range (*r*_0_ selection results in Appendix B).

### 2.6 Sensitivity analysis for entropy

For an understanding of the impact of different hallmarks on the development of ITH in the tumor, we conduct Morris one-at-a-time (MOAT) sensitivity analysis. Since melanocyte proliferation and apoptosis base rates are taken from the literature, we focus on the remaining hallmarks. The driver gene strength *f* is involved in the implementation of all hallmarks. We vary the base mutation rate *r*_0_ for genetic instability, and for immune evasion we look at both the immune base proliferation *p_i_*_0_ and the immune base recruitment rate *s*_0_. For comparative study, we set up both hot and cold tumors to investigate how the density and chemotaxis of CTLs contribute to the impact of the immune evasion hallmark in BEP-HI. A BRAF tumor with immune evasion (IE) served as a cold tumor (*BRAF_IE_*_+_) while BRAF without IE served as a hot tumor (*BRAF_IE_*_−_).

### 2.7 Introduction of BRAF-mutant motility

In the extended BEP-HIM model, on acquiring a mutation in BRAF driver gene, the cell gains the ability to move with a probability of *m*_0_. If there is an empty spot in the neighborhood, the cell moves there. Otherwise, it moves to an occupied spot and pushes all the cells in that direction out by one grid point. To find a realistic range of values for *m*_0_, we compare the invasive metrics of convexity and border fractal dimension for benign and malignant melanoma image dataset (details in Appendix B).

### 2.8 Spatial analysis using cross-pair correlation function

In order to quantify spatial correlations between subclones, immune cells, and normal cells at a distance *r*, we use the cross-pair correlation function (*cPCF*). *cPCF* indicates how target cell types arrange themselves relatively at different distances from the reference type, with *cPCF* = 1 when there is spatial random distribution, *cPCF <* 1 for anti-clustering types and *cPCF >* 1 for clustering types (Bull and Byrne 2023).

## 3 Results

### 3.1 BEP-HI shows emergence of three intratumor heterogeneity modes

As initial validation of the functioning of BEP-HI in producing intratumor heterogeneity (ITH) variation, we investigate how changes in the base mutation rate (*r*_0_) and driver gene strength (*f*, representing the impact of a driver mutation on all cancer hallmarks) influence the emergence of distinct ITH patterns. By systematically varying these two parameters, we see three qualitatively different tumor evolutionary modes emerge in **Figure 2**. We observe clonal sweeps in which a single dominant clone overtakes the tumor due to strong selective advantage. In subclonal mode, multiple subclones coexist and expand, resulting in subclonal sweeps and moderate ITH. Finally, some tumors exhibit fractal heterogeneity with complex spatial structure and high levels of genetic diversity. These results closely mirror the branching evolutionary dynamics previously described (Niida et al. 2015), and provide an interpretable framework for tuning evolutionary pressures in our extended model.

**Fig. 2:**
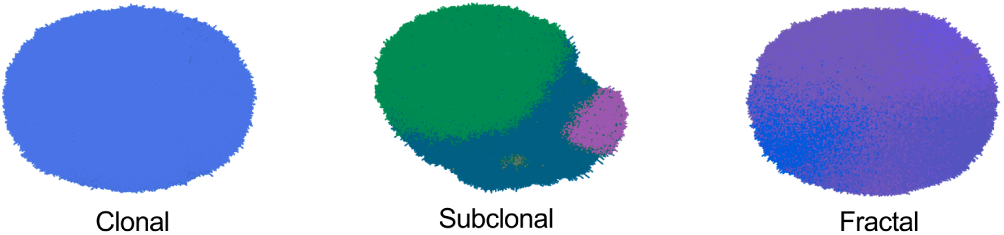
Tumors visualized at the end of our simulations, colored by cell genome, show three distinct ITH modes. We observe (i) clonal sweeps in which a single dominant clone overtakes the tumor; (ii) subclonal mode in which multiple subclones coexist and expand; (iii) fractal heterogeneity with complex spatial structure and high levels of genetic diversity.

### 3.2 Space systematic sampling best captures complex heterogeneity

**Figure 3** compares the accuracy of four sampling strategies across clonal, sub-clonal, and fractal tumors. We plot the error in ITH metrics entropy (*ɛ*) and tumor mutational burden (*β*) for four sampling methods: simple random sampling (SRS), stratified sampling via CLARANS (CC), time systematic (TS), and space systematic (SS). TS consistently underperforms with highest error in both *ɛ* and *β* across all 3 ITH modes, likely due to the stochastic nature of cell mutation and death, which undermines the assumption that temporal order reliably reflects genetic diversity. SRS is adequate for clonal tumors with minimal heterogeneity, but its performance declines in more complex subclonal and fractal regimes. Between the remaining methods, both CLARANS and SS perform comparably for subclonal tumors; however, SS demonstrates superior performance in capturing the spatial complexity of fractal tumors. Furthermore, SS offers substantial computational advantages, averaging 292 seconds per 25,000-cell simulation, compared to 535 seconds for CLARANS. Given the anticipated increase in computational demands for larger populations (e.g., 50,000 cells), spatially systematic sampling is selected as the preferred method for subsequent analyses.

**Fig. 3:**
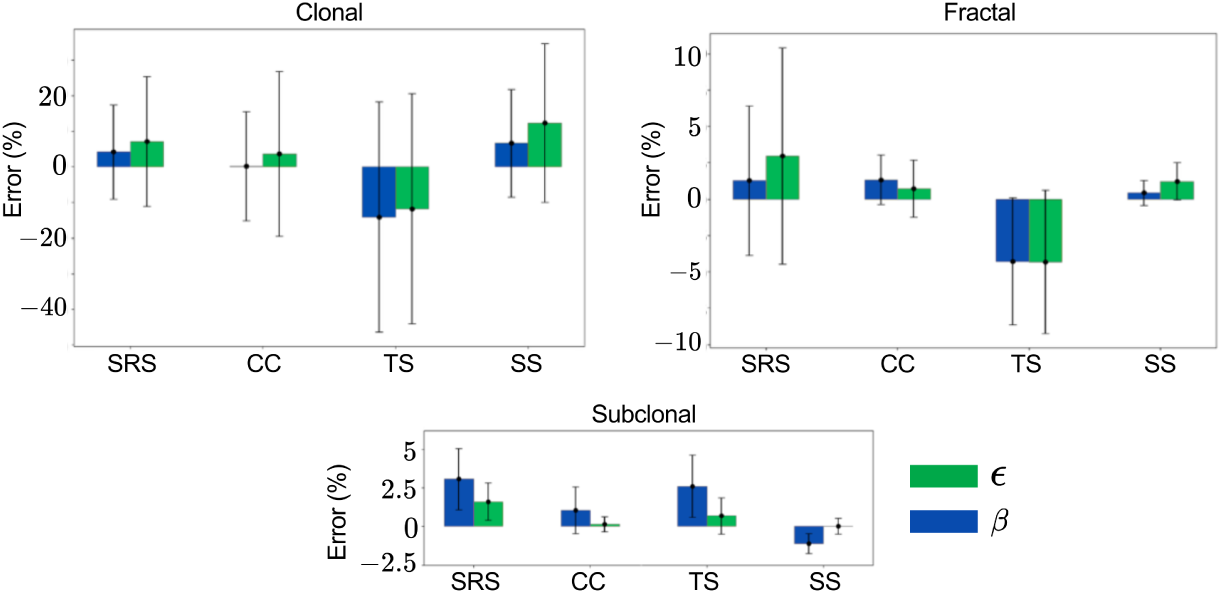
Space systematic is best at capturing spatial heterogeneity. Plots show mean error % in entropy (*ɛ* in green) and average number of mutations (*µ* in blue) in the four different methods: simple random sampling (SRS), stratified sampling via CLARANS (CC), time systematic (TS), and space systematic (SS). The standard deviation is included as error bars.

### 3.3 Driver gene strength governs growth

Driver gene strength is calibrated by matching simulated time to detectable tumor size (*c_max_* = 50, 000 cells) with reported growth times for superficial spreading melanoma. The base mutation rate is selected to produce realistic post-detection growth kinetics. **Figure 4a** plots the time to reach *c_max_* as a function of *f*, showing a monotonic decrease in time with increasing *f* — i.e. stronger drivers yield faster-growing tumors. This places SSM-like growth kinetics in the range *f* ≈ 1.4 − 1.6 (slower growth). Increasing *f* beyond this range drives markedly faster growth, consistent with NRAS- or KIT-associated nodular melanomas that can become detectable within weeks to months.

**Fig. 4:**
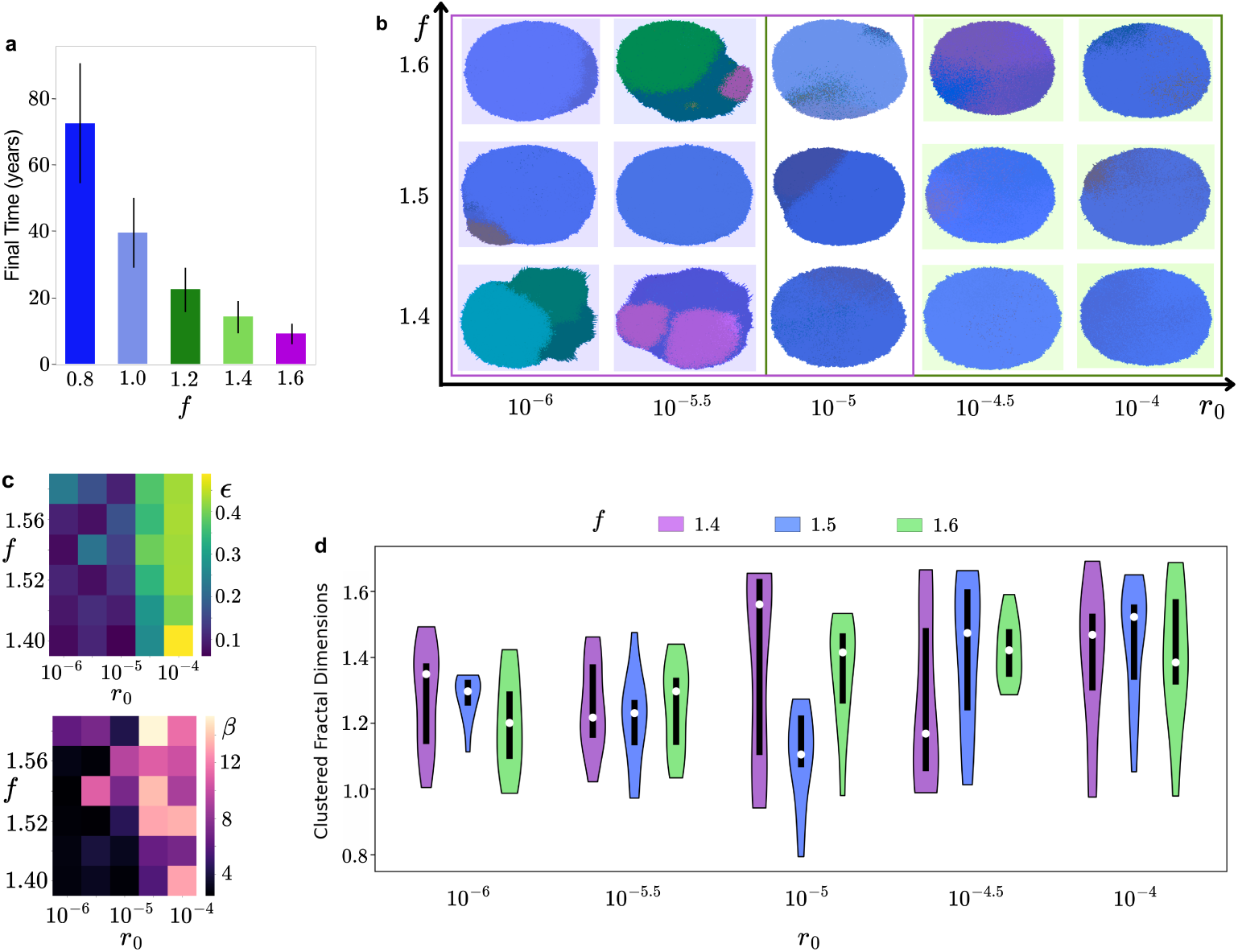
Decoupling of growth kinematics and ITH in tumors. **(a)** Plot of the time taken for simulations to reach at least 50,000 ABM cells against the driver gene strength *f*. **(b)** Examples of tumor visualizations for different *r*_0_*, f* parameters showcasing the heterogeneity modes that emerge with *f* ∈ [1.4, 1.6]. **(c)** Heatmaps of ITH metrics entropy (top panel) and of average number of mutations (bottom panel) with varying *r*_0_ and *f*. **(d)** Average clustered fractal dimension (*FD*) is grouped by *r*_0_ values with the violin plot colors representing different *f* values. The white circles indicate median values.

### 3.4 Mutation rate shapes distinct evolutionary patterns of ITH

For an initial understanding of the ITH dynamics in BEP-HI, we consider distributions of our three quantifying metrics over a (*r*_0_*, f*) parameter space. We observe that intratumor heterogeneity (ITH) is primarily governed by the base mutation rate *r*_0_.

This became evident in the tumor visualizations shown in Figure **Figure 4b**, where we present example tumors generated using our estimated (*r*_0_*, f*) values. We note that for *r*_0_ *<* 10^−5^, simulations produce clonal or subclonal tumors, whereas for *r*_0_ *>* 10^−5^, they shift almost exclusively into high-ITH fractal cases. No comparable transition is observed with changes in *f*. To quantify these observations, we plot two (*f*, *r*_0_) heatmaps in **Figure 4c** where the top panel shows entropy (*ɛ*) and the bottom panel shows mean mutation count (*β*). In both panels, values increase primarily along the *r*_0_ axis, indicating that intratumor heterogeneity (*ɛ*) and mutational burden (*β*) scale with the base mutation rate across all *f*. There is no consistent trend with *f* in either heatmap, underscoring that driver strength governs growth rate, while mutation rate (*r*_0_) governs genetic diversity. This is further reinforced by **Figure 4d** which shows clustered fractal dimension (*FD*) distributions grouped on the x-axis by *r*_0_ and colored by *f* values. *FD* serves as a quantitative measure of space-filling capacity, with higher values indicating more complex morphologies that more efficiently explore and occupy the surrounding tissue. In this context, the fractal dimension can be thought of to reflect the internal space-filling complexity of overlapping or nested subclones rather than external perimeter roughness. *FD* increases with an increase in *r*_0_ confirming that tumors in the high mutation regime corresponding to fractal heterogeneity have significantly higher fractal dimensions. Interestingly, tumors in the fractal regime, despite their high genetic diversity and complex evolutionary trajectories, exhibit smoother, more regular tumor shape than the lower *FD* associated subclonal tumors as seen in Fig **Figure 4b**. These findings underscore the importance of employing multiple spatial metrics to comprehensively capture the phenotypic consequences of intratumor heterogeneity. Together with **Figure 4a**, these plots demonstrate a decoupling of kinetics and diversity: *f* sets the pace to detectable size, whereas *r*_0_ sets the level of ITH observed at that size.

### 3.5 Immune recruitment most strongly influences ITH

Our aim of predicting ITH modes for simulated tumors demanded a deeper study of how the immune system affects ITH in our model. In the entropy targeted Morris sensitivity analysis (MOAT) shown in **Figure 5a**, *µ*^∗^ indicates the degree of influence a parameter has on entropy, and *σ* indicates the non-linearity and interdependence of the parameter. The overall low *µ*^∗^ values for all the parameters show that no one parameter completely controls ITH. Additionally, the high *σ* suggests that the effects of parameters is non-linear and interdependent, especially for *f* which is expected due to its role in changing hallmark action probabilities, and for *s*_0_ which determines strongly the rate at which high antigenicity subclones are killed. Since *s*_0_ has the highest *µ*^∗^ influence index for entropy, the recruitment of immune cells has the most significant influence on the development of ITH. Therefore, we further investigate immune infiltration, including how immune cells enter, navigate, and respond to local cues such as the antigenicity associated ISF gradient.

**Fig. 5:**
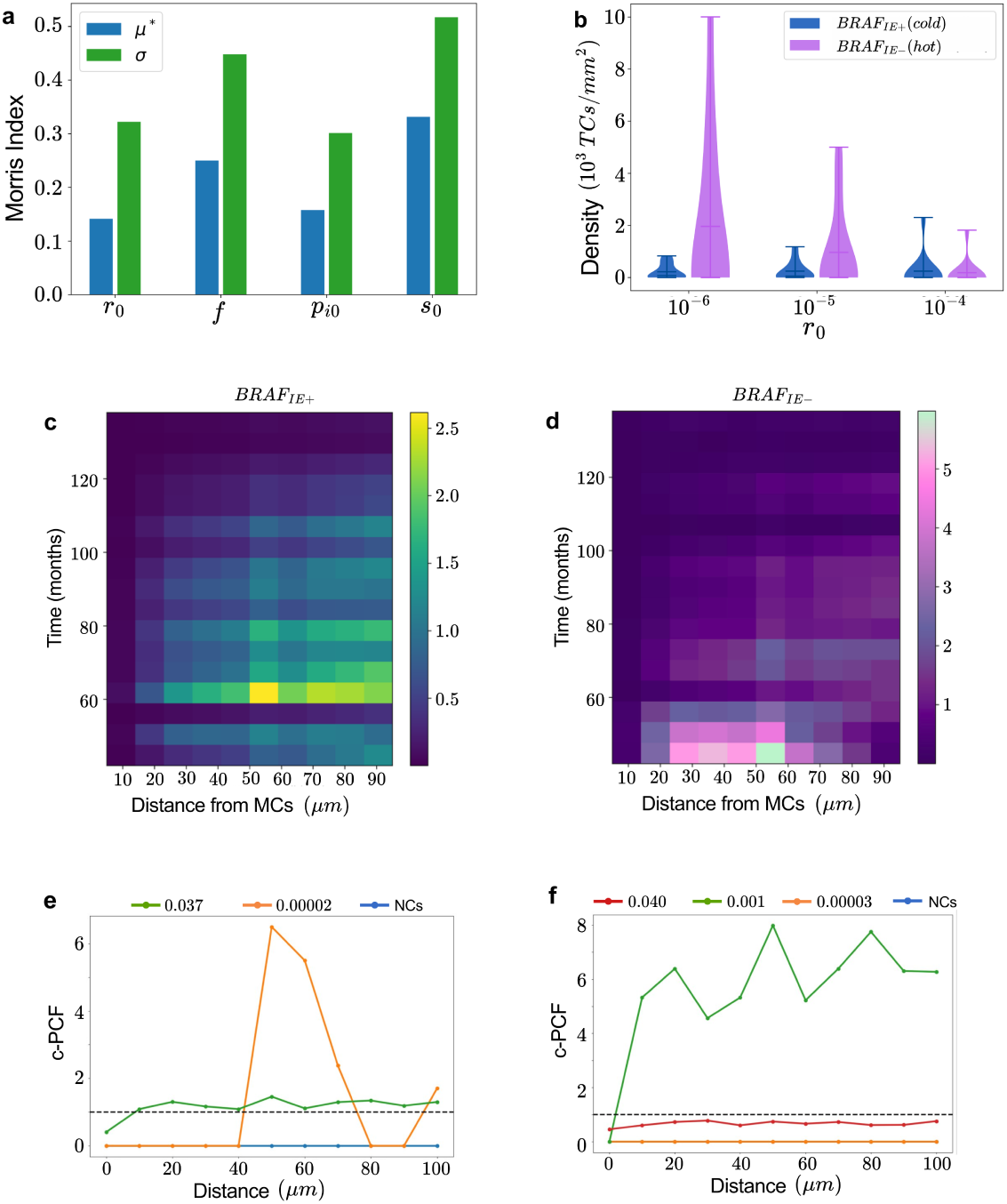
Immune evasion is the most influential hallmark in determining ITH. **(a)** MOAT analysis with entropy as the target. *µ*^∗^ indicates the overall influence of the parameter and *σ* indicates the degree of interaction with other parameters. **(b)** Density of immune cells in (left) cold *BRAF_IE_*_+_ tumor and (right) *BRAF_IE_*_−_ hot tumor at 5 years. **(C)-(D)** Average number of CTLs at a particular distance from a MTCs, normalized tumor cell count in cold tumors in **(c)** and hot tumors in **(d)**. **(e)-(f)** Cross-PCF analysis of how immune cells (reference) arrange themselves around different antigenicity cells. Each line represents a cell type labeled by its antigenicity. **(e)** A typical example of *cPCF* plot where CTLs are strongly clustering with the highest antigenicity cells and **(f)** shows an example exception to this trend.

In **Figure 5b**, we plot the density of CTLs inside the convex hull of the tumor after 5 years for *f* = 1.6 and varying *r*_0_ values, with 10 runs per parameter set, for both BRAF with immune evasion (cold) and BRAF without immune evasion (hot). CTLs are more present in hot tumors than cold and this is most evident for *r*_0_ = 10^−6^. As the base mutation rate *r*_0_ increases, we see an increase in CTL density for (*BRAF_IE_*_+_) but a decrease in CTL density for (*BRAF_IE_*_−_) hot tumors. We know from the TMB heatmap (bottom panel in **Figure 4c)**, that as *r*_0_ increases, there is an associated increase in the tumor mutational burden (*β*), i.e greater number of random mutations per cell. These random mutations could increase antigenicity of the melanoma cell which in turn leads to an increase in ISF-dependent immune cell recruitment and proliferation, as observed for cold tumors. The same mechanism could explain the behavior in hot tumors except that for high *r*_0_ values, the initial sweep of *BRAF_IE_*_−_ cells with high antigenicity leads to earlier recruitment and proliferation of immune cells. The CTLs then target and kill the high antigenicity cells leaving behind a less immunogenic tumor at 5 years.

To explore this theory, we generate heatmaps representing the average number of CTLs at a particular time and distance from MTCs (**Figure 5c** for cold and **Figure 5d** for hot tumors) to gain insight into how CTL behavior changes over time. The CTL count is normalized by the MTC count and there are 10 runs for both the hot and cold cases. CTLs reach their highest density much later in the simulation for cold tumors than for hot tumors, supporting our theory for the low density in hot tumors at 5 years. Additionally, the maximum density in hot tumors is twice that in cold tumors, and occurs at much closer distances indicating that CTLs are more readily recruited and able to track MTCs in hot tumors.

In order to ensure that CTLs are efficiently tracking MTCs in the cold tumor environment, we produce cross-PCF analysis with CTLs as the reference and cells grouped by antigenicity as the targets. The dashed line at *cPCF* = 1 represents normal distribution while *cPCF >* 1 is clustering and *cPCF <* 1 is anti-clustering at a particular distance. **Figure 5e** is an example of the majority of cases where the highest antigenicity cells are clustering at smaller distances indicating efficient MTC tracking by CTLs. However, there are some exceptions that occur - example in **Figure 5f** - where a lower antigenicity cell is much more clustering with the CTLs at all distances. These exceptions could be attributed to cases where CTLs are recruited recently to the tumor periphery and are therefore closer to one cluster of cells. Both these figures also present that normal cells with *a* = 0 remain strongly anti-clustering with CTLs (*cPCF* = 0) at short distances, as in virtually all simulations.

### 3.6 Tumor cell motility drives border irregularity

In order to match simulated tumor morphologies to clinical SSM tumors, and to look for correlations between ITH and morphology, we conduct image analysis on benign and malignant tumors and plot distributions of convexity *C*and border fractal dimension *FD_b_* in **Figure 6a**. There is a large range of *C* values, going as low as 0.1 in the case of malignant tumors. The distribution of *FD_b_* is similar across malignant and benign tumors with a median of about 0.3. We then investigate the morphology of our simulated tumors using these metrics *C* and *FD_b_*, and plot these results under varying motility probabilities *m*_0_ in **Figure 6b**. In BEP-HI (*m*_0_ = 0), while tumor visualizations suggest greater irregularity in subclonal tumors, both *C* (≈ 0.6 − 0.72) and *FD_b_* (≈ 0.98 − 1.08) have a very small range of values which shows that there is no significant difference in morphology between the ITH modes. Specifically, the high *C* values indicate a regular, circular shape of the tumor and low *FD_b_* values indicate that the boundary is smooth.

**Fig. 6:**
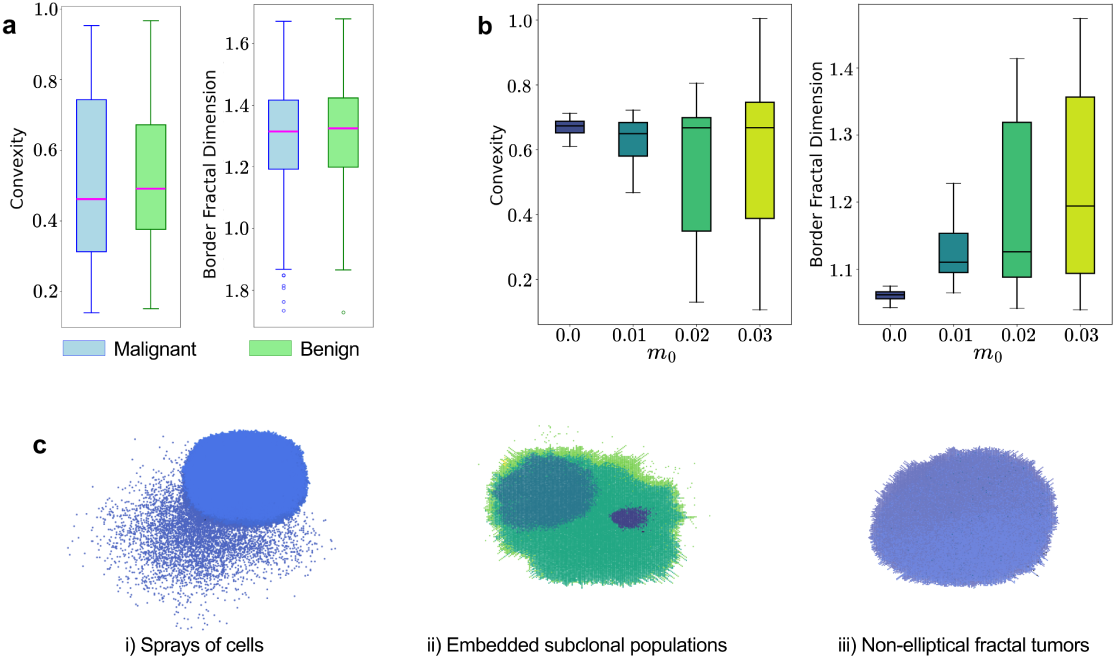
Introduction of motility to BRAF-melanoma cells reveals more irregular morphologies connected to ITH modes. (**a**) Distributions of convexity (*C*) and border fractal dimension (*FD_b_*) from image analysis of malignant and benign melanoma tumors. (**b**) Boxplots of *C* and *FD_b_* distributions for simulated tumors with different motility probabilities *m*_0_. (**c**) Emergence of new morphologies including (i) spray of highly motile cells, (ii) subclonal clusters being embedded within other subclonal clusters, and (iii) non-elliptical fractal tumors.

To reproduce the irregular tumor morphologies observed in clinical examples, we introduce motility *m*0 *>* 0 for the BRAF-mutant melanoma cells, resulting in the BEP-HIM model. After calibration, we find that increasing motility into the range *m*0 ≈ 0.2–0.3 yields convexity and fractal border dimension values comparable to those seen clinically. In the case of convexity values for *m*_0_ *>* 0 in **Figure 6b**, median *C* remains around 0.7 as *m*_0_ increases, and in fact is much higher than the ≈ 0.45 median for *C* in clinical cases. However, we get a much larger range for *C* as *m*_0_ increases, values becoming similar to **Figure 6a**. *FD_b_* shows a much more direct relation with *m*_0_, again matching the range of clinical *FD_b_* when *m*_0_ ≈ 0.2 − 0.3. We visualize some of the tumors produced in BEP-HIM (with *m*_0_ *>* 0) in **Figure 6c**, and observe new morphologies associated with the different ITH modes: (i) one of the subclones (especially in clonal sweeps) is extremely migratory and breaks away from the main tumor body, (ii) one subclone envelops others when previously subclonal populations usually emerged at the edges of the tumor, and (iii) fractal heterogeneity tumors that are not perfectly round as before. All of these more complicated spatial arrangements, and especially the spread of a subclone away from the main tumor body, explain the significant changes observed in *FD_b_* with changes in *m*_0_.

### 3.7 Spatial relations between cell lineages persist with introduction of motility

To establish how genetically distinct subclones spatially arrange themselves within the tumor, we use cross-pair correlation function (*cPCF*) analysis. We categorize cells by the aggressiveness of their genome which is determined by the count of their driver genes. The most aggressive is used as the reference with the remaining as targets (ranked and labeled by their aggressiveness) and the cross-PCF at varying radial distances plots are in **Figure 7a-7c** for *f* = 1.6 and *r*_0_ = 10^−6^, 10^−5^, 10^−4^. The least aggressive are the normal cells with no driver mutations (blue lines). In all cases, these normal cells remained anti-clustering with the most aggressive cells, particularly at close distances. Also, the second most aggressive clone (labeled 2) is usually highly clustering with the most aggressive. This spatial arrangement makes sense since these are usually a single driver mutation apart indicating that the most aggressive is likely progeny of the second most aggressive. We further observe that even with the introduction of motility in BEP-HIM, the cross-PCF plots remain similar (see **Figure 7d-7f**) suggesting that these spatial patterns are preserved regardless of motility and across ITH modes.

**Fig. 7:**
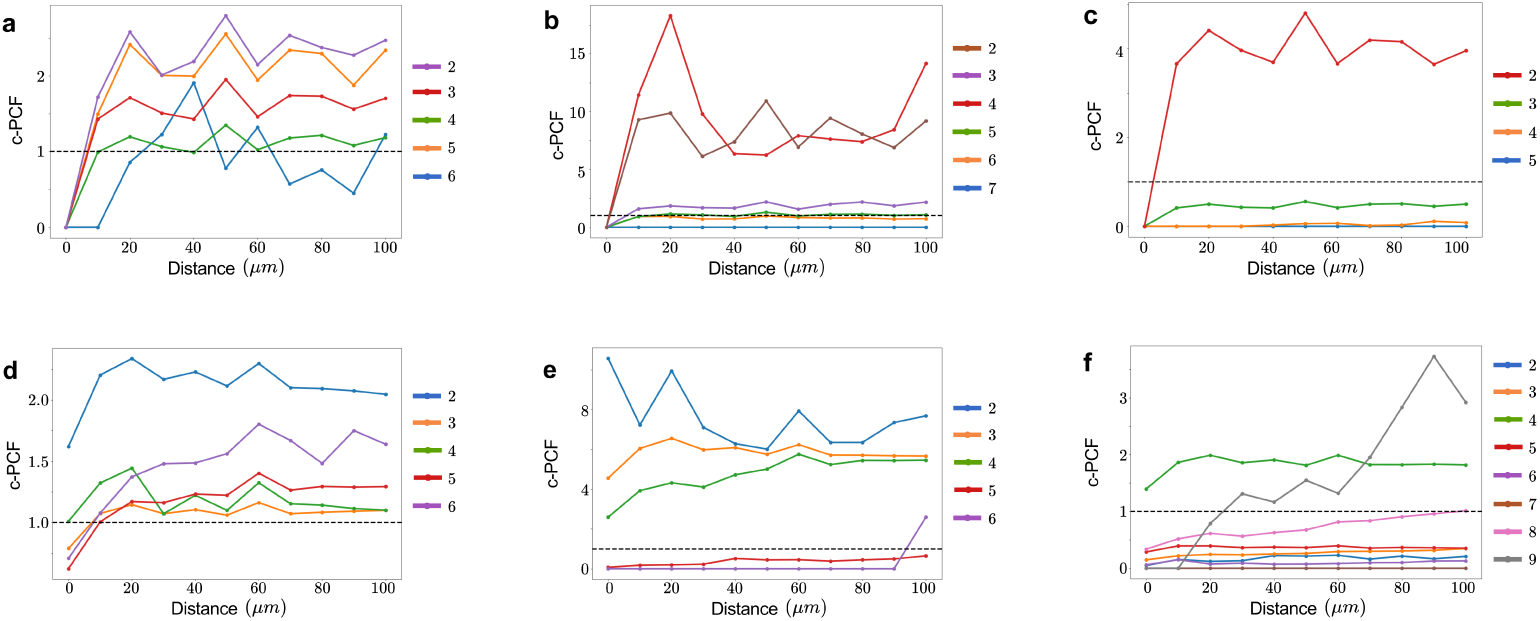
Cross-PCF analysis shows persistent relationship between cell lineages. **(a)-(c)** Plots of cross-PCF against radial distance for BEP-HI with *f* = 1.6 and **(a)** *r*_0_ = 10^−4^, **(b)** *r*_0_ = 10^−5^, and **(c)** *r*_0_ = 10^−6^. The dotted line represents random spatial distribution. We use the most aggressive genome as the reference. The remaining cells are grouped and ranked by aggressiveness and treated as targets with a label of 2 being the second most aggressive and least aggressive being normal cells (blue). **(d)-(f)** cPCF analysis is repeated for BEP-HIM with same *f* and *r*_0_ values as before.

## 4 Discussion

We developed an agent-based model that captures the emergence of intratumoral heterogeneity (ITH) and the spatial growth dynamics characteristic of superficially spreading melanoma (SSM). The model integrates hallmark-specific behaviors and maps them to mutations in seven melanoma-relevant oncogenes, including the SSM-defining founder mutation BRAF. Studies have found a correlation between the presence of CTLs in advanced SSM tumors and clinicopathological characteristics including median tumor size and significantly on median mitotic count (Zablocka et al. 2021). These phenotypic characteristics are connected to several genetic mutations found within the SSM subtype, with 64.3% of SSM cases having BRAF mutations - *BRAF^V^* ^600^*^E^* being most frequently detected (Druskovich et al. 2024). TERT promoters, CDKN2A, and MITF mutations are also common and associated with melanogenesis (melanocyte development and survival) while PTEN is observed in more advanced tumors [(Druskovich et al. 2024), (Guan et al. 2015)]. Furthermore, data from skin cutaneous melanoma - The Cancer Genome Atlas (SKCM-TCGA) can be used to establish any co-occurrence likelihoods of BRAF with the driver genes in BEP-HIM, and to understand how various driver gene combinations may change mutation count, fraction of gene altered, and prognosis. The framework is intentionally designed to be modular, enabling extension to other cancer types by adjusting hallmark behaviors, mutational landscapes, or microenvironmental inputs.

Fractal dimension (*FD*) distinguishes the three ITH modes, showing a clear increase upon transitioning from clonal/subclonal to fractal patterns (**Figure 4b**, **4d**), consistent with clinical use of *FD* to characterize melanoma morphology and aggressiveness. While *FD* ranges vary across prior studies due to methodological and tumor-stage differences [(Piantanelli et al. 2005), (Popecki et al. 2022), (Romero-Morelos et al. 2024)], **Figure 4d** proves the metric is robust in our controlled in silico setting with distributions plotted for 10 simulations per parameter set. *FD* ≈ 1.0 − 1.3 is associated with low ITH and 1.3 − 1.6 with high ITH consistently, with significant variation only around ITH mode transition at *r*_0_ = 10^−5^.

Our analyses reveal a decoupling: driver gene strength *f* determines the time required for a tumor to reach a clinically detectable size, whereas genetic instability *r*_0_ governs transitions in ITH modes. This separation of growth and diversity determinants provides a conceptual tool for understanding why tumors of similar size may exhibit dramatically different evolutionary architectures. We also see a correlation between ITH and tumor mutation burden (*β*), which is an important biomarker in clinical studies. BRAF-mutant melanoma tumors tend to have high TMB, with the SKCM dataset showing a mutation rate of 18 mutations per mega base pairs (Mbp) which is about 10-fold richer than other cancers in TCGA such as glioblastoma multiforme (GBM) (Guan et al. 2015). Additionally, melanoma is associated with low subclonal mutational load (sML), the proportion of subclonal mutations across clusters, which is also an indicator of poor prognosis (Jiang et al. 2024). This supports our conclusion from tuning of *r*_0_*, f* that SSM cases often fall into fractal heterogeneity associated with high ITH. High intra-tumor heterogeneity leads to inefficient recognition of antigens by the immune system due to diluted frequency and spatial heterogeneity, producing tumors that respond poorly to ICI therapy (Craig et al. 2022).

The predominant influence of immune recruitment rate *s*_0_ likely arises because *s*_0_ controls the fundamental timing and magnitude of immune editing pressure. Because recruitment increases with peripheral inflammatory signals, *s*_0_ participates in a feed-back loop that prunes high-antigenicity clones and accelerates the emergence of immune-evasive lineages. Consequently, small perturbations in *s*_0_ can shift the system across three regimes: weak editing (favoring clonal sweeps), intermediate editing (sustaining coexistence of multiple subclones), and strong editing (driving rapid immune escape). These threshold and time-dependent interactions, also visible in the hot vs. cold tumor patterns of immune density over time (**Figure 5c-5d**), are potential reasons why *s*_0_ also strongly influences entropy. Our work supports earlier findings that the evolution and prognosis of melanoma is more heavily influenced by interaction between immune cells and tumor cells than the genetic subtype of driver mutations (Yang et al. 2023). Spatial transcriptomics has shown that T cells infiltrate tumors in a spatially diverse manner, congregating in craters, often found at the margins of melanoma (Ludin et al. 2024) and strongly shaped by cytokine gradients (Centofanti et al. 2023). Our model accounts for these factors of CTL recruitment based on melanoma type through *s*_0_ and on cytokine gradients through a Hill function dependent on immune stimulatory factor (ISF). In particular, *s*_0_ can be informed through immunohistochemistry studies quantifying T cell infiltration in SSM, such as low or high grades, Clark’s method, or density measurements (Zablocka et al. 2021). However, the high interdependence of *s*_0_ with other parameters complicates targeted modulation. By contrast, *r*_0_ has weaker interactions, making genetic instability a more direct and tunable lever for exploring diversity outcomes.

Introducing motility in mutant BRAF cells (*m*_0_ *>* 0) produces border convexity and fractal dimension distributions that better match clinical observations, and also alters internal spatial organization in different ITH modes. These findings agree with our understanding of how cell motility, along with loss of cell adhesion, drives irregular border formation by enabling migration at both the individual and collective level into surrounding tissues in an uncoordinated manner (Ju et al. 2018). While cross-PCF analysis confirms short-range clustering of aggressive subtypes regardless of motility, it is unable to distinguish BEP-HI from BEP-HIM, suggesting that richer spatiotemporal methods—such as persistent homology—are needed to fully characterize dynamic morphological complexity.

Although the model is designed to be flexible, a few limitations constrain its adaptability. Most notably, the current implementation is two-dimensional, which is not well suited to more aggressive melanoma subtypes, such as nodular melanoma, which extend vertically into deeper tissues. However, in our study of a radially growing SSM tumor, a two-dimensional model enables a tractable exploration of growth and genetic diversity. Similarly, our treatment of the immune microenvironment is simplified and does not yet incorporate stromal heterogeneity, vasculature, or mechanical constraints. Recent developments in 3D volumetric imaging has made it possible to have realistic spatio-phenotypic and dynamic tumor traits in 3D models (van Ineveld et al. 2022). Tracking of individual cell fates and greater resolution in these processes could be used to capture the entire cellular heterogeneity of cancer and produce datasets to tune stromal or vascular models [(van Ineveld et al. 2022), (Yapp et al. 2025)]. This simplification of TME dynamics allows for a much clearer relationship to be established between specific immune actions and ITH which will be crucial in future work with optimizing the administration of immunotherapy to control for ITH in SSM tumors. Furthermore, any additions would be computationally taxing, necessitating further restrictions on analysis techniques such as entropy calculations for ITH estimation which is already limited to a sample size of 1/10th of the population.

Taken together, our BEP-HIM framework captures key genetic, spatial, and morphological features of superficially spreading melanoma while establishing mechanistic links between cancer hallmarks, evolutionary dynamics, and tumor architecture. A major novelty of the model lies in its integration of specific driver genes, explicit immune-editing feedback, and motility-dependent morphological evolution within a single spatially resolved agent-based platform. This combination enables biologically grounded exploration of how genetic instability, immune pressure, and phenotype switching interact to shape ITH and treatment-relevant trajectories. By allowing mutation processes to map directly onto real melanoma driver genes, the model provides a foundation for generating virtual tumor cohorts that can be parameterized using sequencing, spatial transcriptomics, and imaging data. Such virtual cohorts can be used to test hypotheses about the emergence of subclonal therapeutic resistance, identify early-warning signatures of immune escape, evaluate the impact of targeted or combination therapies, and design sampling strategies that better capture clinically meaningful heterogeneity. Ultimately, the BEP-HIM model offers a flexible and extensible platform for connecting evolutionary theory with experimental and clinical melanoma research, and for guiding the development of personalized treatment strategies that explicitly account for spatial and genetic diversity.

## Statements and declarations

The authors declare that they have no known competing financial interests or personal relationships that could have appeared to influence the work reported in this paper.

## Data availability

All original data supporting the findings of this study is available within the paper. A publicly available Kaggle dataset of melanoma images used during calibration of motility is available at: https://www.kaggle.com/datasets/hasnainjaved/melanoma-skin-cancer-dataset-of-10000-images?. Further inquiries can be directed to the corresponding author.

## Appendix A

### A.1 Melanocyte properties

Each melanocyte cell has a genome consisting of 50 genes of which 7 are driver genes associated with melanoma. These driver genes are listed in **Table 1**, with the hallmark actions which they affect probabilities for, taken from existing literature.

We add to the melanocyte cells an antigenicity property *a*, which is *a* = 0 for normal cells. In the case of tumor cells, the base antigenicity *a*_0_ is randomly chosen so that *a*_0_ ∈ [1, 2]. Then *a* = *a*_0_10*^f^*^(^*^dc^*^°−^*^dc^*^)^, to account for driver genes that help in immune evasion reducing antigenicity while the rest of the drivers increase antigenicity. The antigenicity of a tumor cell determines its contribution to the background ISF which is critical in establishing how CTLs behave in relation to MTCs.

### A.2 Melanocyte updates

At each time step, melanocytes have the probabilistic opportunity to undergo proliferation and apoptosis. If a cell is marked for proliferation, only then do its wild genes have the independent option to mutate. Additionally, division action is applied to marked cells in a random order. The new cell is placed randomly in an empty space in the Moore neighborhood of the original cell if it is available. Otherwise, one of the eight directions is chosen by taking probability for that direction to be the inverse of the distance to the next empty spot in that direction (Niida et al. 2015). Then all cells in that direction are moved out so that the new cell is still placed in the Moore neighborhood of its parent. All action probabilities for melanoma are included in Table 2.

**Table 2:**
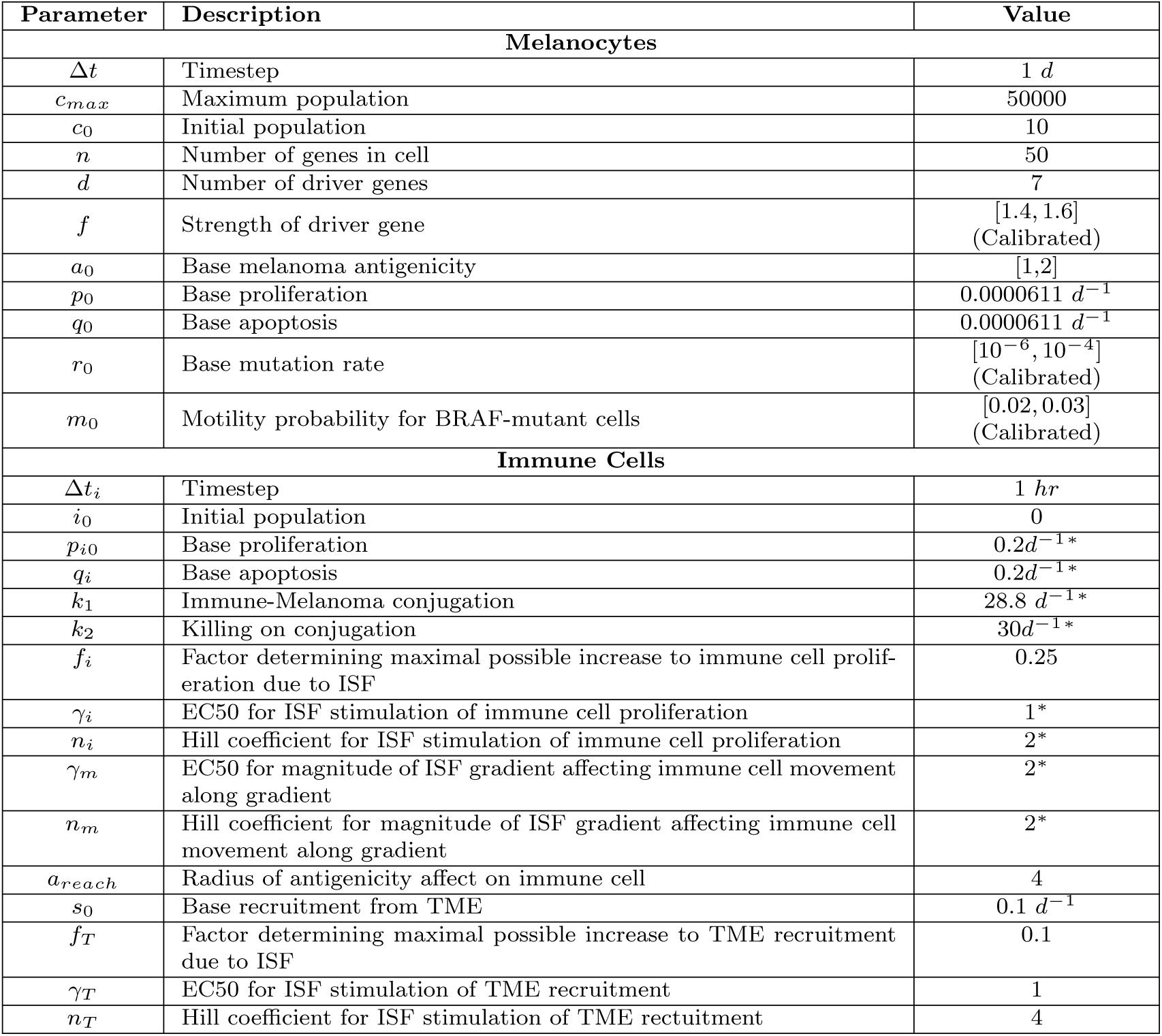
ABM Parameters. ^∗^ refers to values that have been taken from (Bergman et al. 2024). There is very little information about melanocyte base proliferation in literature so we started with a balanced state where *p*_0_ = *q*_0_. During each time step, any cell action happens with a probability P computed using the exponential distribution with rate parameter R such that P = 1 − *e*^−RΔ^*^t^*.

| Parameter | Description | Value |
| --- | --- | --- |
| **Melanocytes** |  |  |
| $\Delta t$ | Timestep | 1 *d* |
| $c_{max}$ | Maximum population | 50000 |
| $c_0$ | Initial population | 10 |
| $n$ | Number of genes in cell | 50 |
| $d$ | Number of driver genes | 7 |
| $f$ | Strength of driver gene | [1.4, 1.6]<br>(Calibrated) |
| $a_0$ | Base melanoma antigenicity | [1, 2] |
| $p_0$ | Base proliferation | 0.0000611 $d^{-1}$ |
| $q_0$ | Base apoptosis | 0.0000611 $d^{-1}$ |
| $r_0$ | Base mutation rate | $[10^{-6}, 10^{-4}]$<br>(Calibrated) |
| $m_0$ | Motility probability for BRAF-mutant cells | [0.02, 0.03]<br>(Calibrated) |
| **Immune Cells** |  |  |
| $\Delta t_i$ | Timestep | 1 *hr* |
| $i_0$ | Initial population | 0 |
| $p_{i0}$ | Base proliferation | $0.2d^{-1*}$ |
| $q_i$ | Base apoptosis | $0.2d^{-1*}$ |
| $k_1$ | Immune-Melanoma conjugation | $28.8 d^{-1*}$ |
| $k_2$ | Killing on conjugation | $30d^{-1*}$ |
| $f_i$ | Factor determining maximal possible increase to immune cell proliferation due to ISF | 0.25 |
| $\gamma_i$ | EC50 for ISF stimulation of immune cell proliferation | 1* |
| $n_i$ | Hill coefficient for ISF stimulation of immune cell proliferation | 2* |
| $\gamma_m$ | EC50 for magnitude of ISF gradient affecting immune cell movement along gradient | 2* |
| $n_m$ | Hill coefficient for magnitude of ISF gradient affecting immune cell movement along gradient | 2* |
| $a_{reach}$ | Radius of antigenicity affect on immune cell | 4 |
| $s_0$ | Base recruitment from TME | $0.1 d^{-1}$ |
| $f_T$ | Factor determining maximal possible increase to TME recruitment due to ISF | 0.1 |
| $\gamma_T$ | EC50 for ISF stimulation of TME recruitment | 1 |
| $n_T$ | Hill coefficient for ISF stimulation of TME recruitment | 4 |

### A.3 Cytotoxic T lymphocytes updates

To model the enhanced immune evasion hallmark, we introduce an immune cell, cytotoxic T lymphocyte (CTL), as an agent that has a base proliferation probability *p_i_*_0_, constant death probability *q_i_*, and can conjugate with a tumor cell to kill it with a probability *k_i_*. The immune cells are also recruited from the TME to the periphery of the tumor at a rate influenced by the antigenicity of the cell at the periphery at any time step. Unlike the static tumor cells, immune cells are capable of moving to the tumor cells by chemotaxis down an immune stimulatory factor (ISF) gradient. This factor performs a role similar to CXCL9 cytokine and its concentration is determined by the antigenicity of local cells and determines the interactions between MTCs and CTLs (Bergman et al. 2024).

The probability of an immune cell killing a tumor cell is also affected by the antigenicity of the tumor cell and is calculated as *k_i_* = *k*_1_*k*_2_10^−^*^fdc^*. The relevant parameter values can be found in Table 2.

We assume these contributions are local, that is they are maximized near the tumor cell and decay to 0 far from the tumor cell. To model this, we use a Gaussian function centered at the tumor cell with covariance given by the identity matrix where the units are one cell length. These functions are then scaled based on the antigenicity of the tumor cell giving *F* which represents the ISF concentration at a given point *h* in the TME and letting a(x) be the antigenicity of tumor cell x located at L(x):

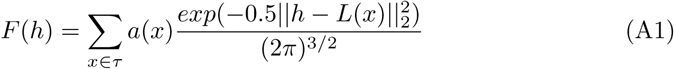

*τ* represents the set of tumor cells within a radius of 4 from *h* for computational simplicity and because any tumor cells further than this from *h* would have insignificant contribution to the ISF concentration at *h*. For an immune cell *y*, the ISF concentration at its location is given by *F* (*y*). When an immune cell moves, it first samples the local ISF gradient, *G*(*y*) ≡ ∇*F* |*_y_*, converting the norm of this vector into a bias

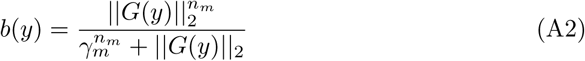

where *n_m_* = 2 is the Hill coefficient for magnitude of ISF gradient affecting immune cell movement along gradient and *γ_m_* = 2 is the EC50 for magnitude of ISF gradient affecting immune cell movement along gradient. Additionally, the proliferation rate is individualized to each immune cell as it is scaled by the ISF concentration at its location (*F*) using a Hill function.

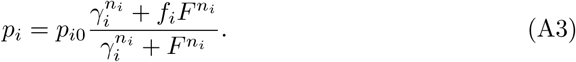

### A.4 Model initialization

The model is initialized with 10 melanocytes of which only 1 randomly selected cell acquires a mutation in the BRAF driver gene and becomes a melanoma cell with non-zero antigenicity. All melanocytes are placed together at the start. There are no immune cells initialized, they must be recruited from the TME.

### A.5 Model parameters

All model parameters for both the melanocyte cells and the immune cells can be found in Table 2. The immune cell parameter values marked with ∗ are from work on an agent-based model for the immune landscape in bladder cancer (Bergman et al. 2024). The remaining immune parameters are estimated to control for immune cell presence in the tumor and allow for the tumor to reach a detectable size. The parameters for driver gene strength *f*, and BRAF-melanoma motility *m*_0_ are calibrated in the main text while selection of realistic mutation rates *r*_0_ is shown in Appendix B. Time steps for agents are separated to account for the different cell cycle timescales for normal melanocytes and immune cells, along with computational limitations.

## Appendix B

### B.1 Visual analaysis of ITH

We use a similar strategy to Niida et. al, and sample *m* cells (10% of the final population) to obtain an *n*x*m* mutation profile matrix **M**, where each of the *m* columns is the *n*-dimensional genome vector **g** for each cell (Niida et al. 2015). Then we use principle component analysis on **M** to produce the first three principle components (PCs) which we use as RGB color samples. Then we multiply **g** for each cell with each of the RGB PCs and sum these to produce a unique coloring for each **g**.

### B.2 ITH Sampling Strategies

To improve upon the limitations of SRS—particularly in simulations where localized subclonal clusters emerged—we implemented stratified sampling based on the genetic states of the cells. This was operationalized using the CLARANS clustering algorithm, allowing us to perform proportional sampling within genetically distinct clusters. In addition, we assessed two systematic sampling approaches. The first, termed systematic spatial sampling (SS), involved overlaying a regular grid on the tumor domain and sampling genetic data from cells located at fixed spatial intervals. The second, systematic temporal sampling (TS), exploited the inherent ordering of cell creation in the BEP-HI model. Because cells were appended to a list upon division and removed upon death, sampling every tenth entry provided a temporal cross-section of cell ages.

### B.3 Growth Rate Analysis

We look at the average diameter, calculated as the mean of the length of the long and short axis. Diameters are measured at the end of the simulations for each parameter set to consider which parameters get us closer to the reality of melanoma growth features. We compare the growth rate of superficially spreading melanoma (SSM) after detection, which ranges between 0.08 to 0.76 mm/month dependent on the mitotic rate (Liu et al. 2006), with the diameter in the last three months of simulation (**Figure 8**) when the simulated tumor is close to detectable size. It seems that the simulations with both low drive gene strength *f* and base mutation *r*_0_ have a much more linear looking growth that is significantly slower indicating that these may not be the most useful parameters for modeling real SSM.

**Fig. 8:**
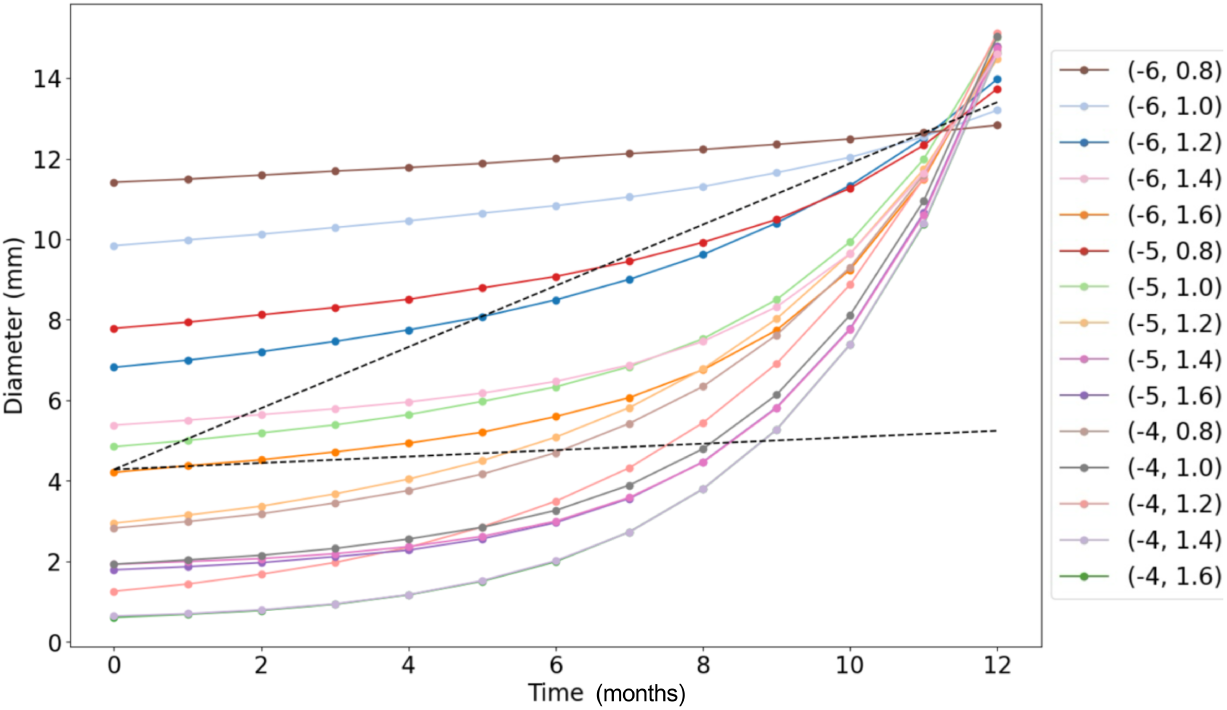
The diameter in mm of the tumor, with the assumption of a cell being a square of side 0.01 mm. Time refers to the last 12 months of each simulations. For each parameter set, results are averaged over 20 simulations. The dashed lines display the range of literature values for melanoma growth rate, and with a scaling of 1:5 for ABM to real cells, we get simulation growth rates within this range.

### B.4 Calibrating Cell Motility

For comparison of morphological features, we use as reference a dataset of 10, 000 benign and malignant melanoma lesions available through Kaggle. The images are processed as shown in **Figure 9**, where parameters for the fuzzing and defuzzing are adjusted to account for factors such as skin texture and contrast of melanoma to skin color (Ali et al. 2020). Additionally, we include removal of hair, rulers, and perfectly circular features which are often the microscope lens in the image. This produces a binary mask, and boundary of this mask is extracted for the calculation of boundary fractal dimension (*FD_b_*) using the box counting method. We also get the length of the largest contour in the mask and find the perimeter and convex hull for convexity calculations.

**Fig. 9:**
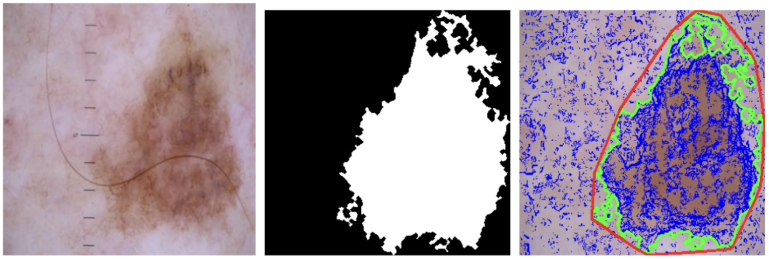
Image processing on melanoma dataset: (i) original image, (ii) binary mask, (iii) overlay with convex hull in red and boundary in green.

### B.5 MOAT Sensitivity Analysis

Morris sensitivity analysis is implemented in Python SALib to generate a set of simulations with 5 trajectories and 4 runs per parameter set to be analyzed. In addition to entropy targeted analysis for BEP-HI, **Figure 10a** shows how cluster fractal dimension metric (FD) for ITH is affected by the four parameters: base mutation rate (*r*_0_), driver gene strength (*f*), base immune proliferation (*p_i_*_0_), and immune recruitment (*s*_0_). The general results match **Figure 5a**, with *s*_0_ being the most influential with non-linear interactions. Furthermore, **Figure 10b** shows that these relationships persist in BEP-HIM with motility being less influential to ITH (entropy) than immune evasion parameters.

**Fig. 10:**
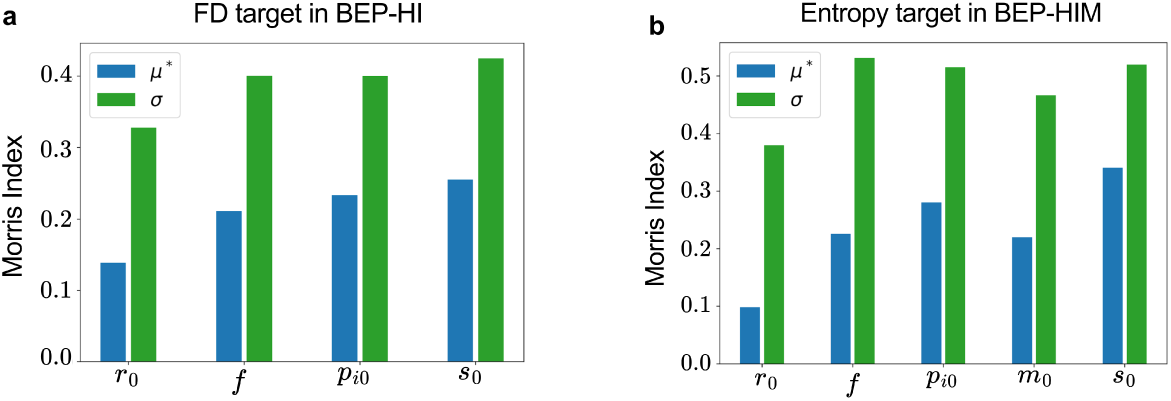
Additional sensitivity analyses support *s*_0_ as most influential to ITH. Plots of Morris sensitivity indices for **(A)** BEP-HI with fractal dimension as the metric for ITH, and for **(B)** BEP-HIM with entropy as ITH metric.

## Notes

### Competing Interest Statement

The authors have declared no competing interest.

